# Scarless gene tagging of transcriptionally silent genes in hiPSCs to visualize cardiomyocyte sarcomeres in live cells

**DOI:** 10.1101/342881

**Authors:** Brock Roberts, Joy Arakaki, Kaytlyn A. Gerbin, Haseeb Malik, Angelique Nelson, Melissa C. Hendershott, Caroline Hookway, Susan A. Ludmann, Irina A. Mueller, Ruian Yang, Susanne M. Rafelski, Ruwanthi N. Gunawardane

## Abstract

We describe a multi-step CRISPR/Cas9 gene editing method to create endogenously tagged GFP-fusions of transcriptionally silent genes in human induced pluripotent stem cells (hiPSCs), allowing visualization of proteins that are only expressed upon differentiation. To do this, we designed a donor template containing the monomeric EGFP (mEGFP) fusion tag and an mCherry selection cassette delivered in tandem to a target locus via homology directed repair (HDR). mCherry expression was driven by a constitutive promoter and served as a drug-free, excisable selection marker. Following selection, the mCherry cassette was excised with Cas9, creating an mEGFP-fusion with the target gene. We achieved scarless excision by using repetitive sequences to guide microhomology-mediated end joining (MMEJ) and introduce linker sequences between the mEGFP tag and the target gene. Using this strategy, we successfully tagged genes encoding the cardiomyocyte sarcomeric proteins troponin I *(TNNI1)*, alpha-actinin *(ACTN2)*, titin *(TTN)*, myosin light chain 2a *(MYL7)*, and myosin light chain 2v *(MYL2)* with mEGFP in undifferentiated hiPSCs. This methodology provides a general strategy for scarlessly introducing tags to transcriptionally silent loci in hiPSCs.

## Introduction

Genome editing has revolutionized cell biology with its ability to precisely edit and engineer genes of interest at their endogenous loci (Doyon et al. 2011, Dambournet et al. 2014, Grassart et al. 2014, Mahen et al. 2014, Otsuka et al. 2016, Roberts et al. 2017). Editing human induced pluripotent stem cells (hiPSCs) provides a particularly powerful model for interrogating cellular organization and dynamics in a diploid, non-transformed and relatively stable genomic context. The ability to differentiate gene edited hiPSCs into multiple lineages makes them an ideal model system for disease modeling and regenerative medicine (Drubin and Hyman 2017).

We have previously described a robust method for fluorescent tagging of select proteins at endogenous levels in hiPSCs (Roberts et al. 2017). The precise addition of a fluorescent tag sequence to the host cell genome is central to this approach and is commonly accomplished via homology driven repair (HDR). Since HDR is an inefficient but essential step in this process, a selection strategy must be used to enrich the small population of edited cells. This is often accomplished by drug selection or by flow cytometry, which rely on successful HDR as well as the expression of the tagged fusion protein. Therefore, these approaches cannot be used to enrich for edited cells where the target gene is silent in hiPSCs but expressed upon differentiation. While editing differentiated cells rather than hiPSCs potentially bypasses this hurdle, the proliferative capacity and pluripotency of the undifferentiated hiPSCs make them a simpler and more scalable editing platform with broader downstream applications. Unlike terminally differentiated cells, an edited hiPSC clonal line can be more easily subjected to expansion and extensive quality control, differentiated into multiple lineages and made available as a shared resource. Therefore, an editing strategy is required to endogenously tag genes that are only expressed in differentiated cell types and transcriptionally silent in hiPSCs.

One strategy for selecting cells edited at a silent locus utilizes HDR-mediated delivery of a selection marker (a drug resistance and/or fluorescent protein) under the control of a constitutive promoter. After selection of the edited cells, this sequence is then removed by recombination, most commonly using the Cre/Lox system. This Cre/Lox recombination event results in a 34 base pair residual LoxP scar, which could disrupt endogenous sequences important for proper regulation of the targeted gene (Skarnes et al. 2011).

To overcome the limitations associated with recombinases and transposases, we developed a multi-step editing strategy using CRISPR/Cas9 to add an mEGFP tag to transcriptionally inactive genes in hiPSCs with drug-free selection and a scarless fusion product (Fig. 1). In the first step, an mEGFP tag is delivered via HDR in tandem with an excisable cassette expressing a second fluorescent protein (mCherry) under the control of a constitutive promoter to facilitate enrichment of edited cells. In a second step, the selection cassette is excised with Cas9. We also included repeat rich sequences in the donor template that guides excision via the microhomology-mediated end joining (MMEJ) pathway. As a result, the excision site is deleted, and a customizable in-frame linker is introduced between the endogenous coding sequence and the mEGFP tag.

**Figure 1.**
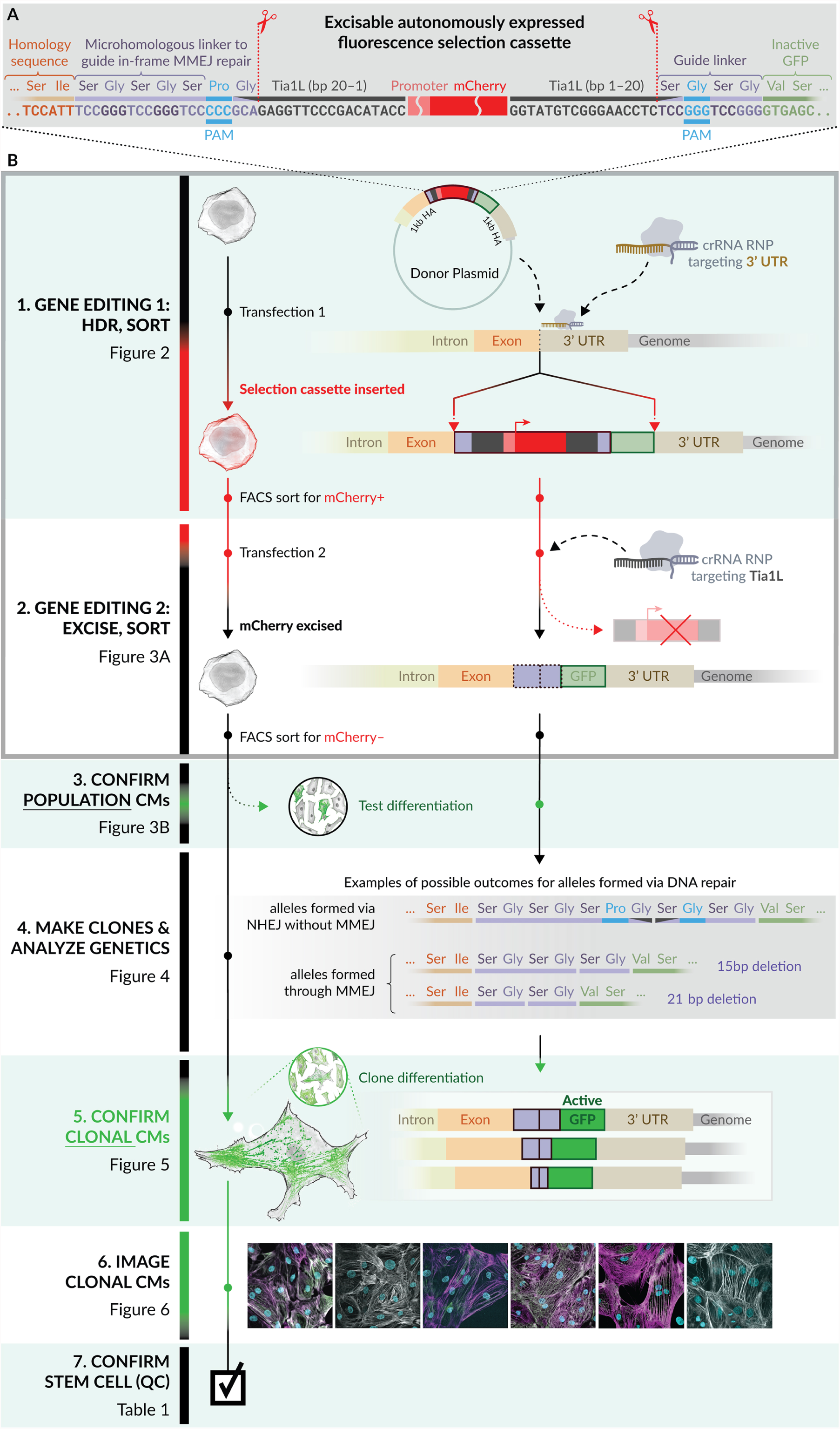
Schematic of multi-step CRISPR/Cas9 mediated targeting via HDR and subsequent microhomology guided excision of the constitutively expressed selection cassette. **A.** Sequences within the TTN donor plasmid (chosen as an example) designed to facilitate scarless MMEJ-mediated excision of the mCherry expression cassette are shown. Promoter and mCherry sequences are not to scale. Tia1L protospacer sequences are in reversed, “PAM-out” orientation. Scissor icons indicate the position of anticipated Cas9 cleavage. **B.** The multistep workflow for editing transcriptionally silent loci and validating success at each step is shown in schematic form. The manuscript figure/table accompanying each step in the workflow is indicated in the left column.

In this report, we demonstrate scarless introduction via this multi-step approach of an in-frame mEGFP tag to the coding sequence of all of the five cardiac genes we targeted, all of which are transcriptionally silent in hiPSCs. The resulting tagged hiPSC lines, after initial targeting, subsequent scarless excision, and validation according to stem cell quality control criteria, represent key structural elements of the cardiomyocyte sarcomere: the Z-disc (*ACTN2*), thin filament (*TNNI1*), thick filaments specific to early atrial and later ventricular developmental fates (*MYL7*, *MYL2*) and the sarcomere spanning protein titin (*TTN*). We generated clonal lines and performed genomic screening to identify monoallelic mEGFP tags with precisely guided in-frame linker sequences lacking a genomic scar. The gene-edited hiPSC clones from all five gene targeting experiments robustly differentiated into cardiomyocytes and demonstrated proper mEGFP localization to the intended sarcomeric structures.

## Results

We developed a multistep gene editing strategy to endogenously tag key cardiac sarcomeric proteins with mEGFP (Fig. 1). To test this strategy we attempted to generate clonal gene edited hiPSC lines for five genes expressed specifically in cardiomyocytes: *TNNI1,* encoding the myofibril contractile regulator slow skeletal Troponin I1; *ACTN2,* encoding the cardiomyocyte-specific actin regulator alpha actinin 2; *TTN*, encoding to the sarcomere spanning structural protein titin; and *MYL7* and *MYL2*, which respectively encode myosin motor proteins expressed earlier and in atrial subtypes (MLC2a) and later and in ventricular subtypes (MLC2v) during cardiomyocyte differentiation. We chose the episomally derived and previously characterized WTC-11 (WTC) line as our parental line (Kreitzer et al. 2013). Population RNA-seq of WTC hiPSCs confirmed that these five genes are transcriptionally silent in pluripotent cells and activated during cardiomyocyte differentiation (hiPSC expression data available at allencell.org). These five sarcomeric proteins span a range of expression levels in hiPSC-derived cardiomyocytes, known localization patterns in the sarcomere, and unique developmental expression kinetics for testing the effectiveness of our editing approach.

### Design of the donor template for silent editing and selection

We designed five donor plasmids, each targeting one of the five chosen cardiomyocyte-specific genes, with a selection cassette in tandem with mEGFP. Each plasmid contained 1 kb locus-specific homology regions for each gene and several key features to enable the multi-step editing strategy (Fig. 1B, Step 1).

The first important feature was a fluorescence (mCherry) selection cassette driven by a constitutive promoter. This selection cassette was adjacent to a second downstream fluorescent tag (mEGFP), intended to ultimately be fused to the C-terminus of the gene of interest. Successful donor sequence incorporation via HDR in hiPSCs transfected with Cas9 and target-specific crRNAs resulted in mCherry expression, which served as a surrogate for editing success at these transcriptionally silent loci and allowed for enrichment of putatively edited cells.

A second important feature was the inverted Tia1L protospacer sites flanking the mCherry donor selection cassette to enable excision (Fig. 1A). These protospacers were included to enable CRISPR/Cas9-mediated excision of the selection cassette after mCherry-expressing cells were initially enriched (Fig. 1B, Step 2). The Tia1L target sequence is absent from the human genome and has been used to ligate distinct double strand breaks introduced by Cas9 (Lackner et al. 2015). We designed these sites in the “PAM-out” orientation such that non-homologous end joining (NHEJ)-mediated double strand repair following Cas9 activity would result in an in-frame mEGFP fusion with the target gene (Fig. 1A). The peptide linker sequences incorporated within the Tia1L sites were designed and oriented such that NHEJ-based repair after excision would result in an in-frame coding sequence with 12 bp of residual sequence (encoding Ser-Gly-Pro-Gly) that served as a canonical linker between the mEGFP and the target gene (Fig. 1B, Step 4).

The third feature of the donor template consisted of microhomology-containing sequences composed of hexa- and tri-nucleotide repeats to encode common peptide linkers in the mEGFP-fusion. We hypothesized that MMEJ events utilizing these repeat sequences would bias excision repair outcomes and efficiently delete the residual sequence remaining from Cas9 cleavage and lead to a more predictable and designable linker sequence (Fig. 1A).

### Step 1 - HDR-mediated delivery of the mCherry selection cassette and mEGFP to target loci

To test the feasibility of this editing strategy, we introduced the donor plasmids into WTC hiPSCs using our previously described CRISPR/Cas9 ribonuclear protein (RNP)-mediated electroporation protocol (Roberts et al. 2017) and evaluated the rate of HDR as indicated by the fraction of mCherry-expressing cells with flow cytometry (Fig. 1B, Step 1). The significant increase in mCherry-expressing cells with gene-specific crRNAs compared to mock control transfections with the plasmid alone indicated HDR-mediated incorporation of this large donor sequence at all five loci (Fig. 2 A,B). Because artificial promoters often display variable responses in pluripotent stem cells, we tested versions of the donor cassette with two different promoters: the low expressing hPGK with the first two genes attempted (*TNNI1* and *ACTN2*) and the higher expressing CAGGS for the subsequent genes tested. Since these experiments were conducted sequentially, we did not test each target gene with both promoters. mCherry-positive cells in the *TNNI1* and *ACTN2* experiments (Fig. 2A) displayed modest fluorescence intensity as expected with the hPGK promoter (Fig. 2A,B). Even though the mCherry-positive population was smaller, the CAGGS-driven mCherry fluorescence intensity was comparatively higher and led to greater ease of sorting (Fig. 2A, y-axis).

**Figure 2.**
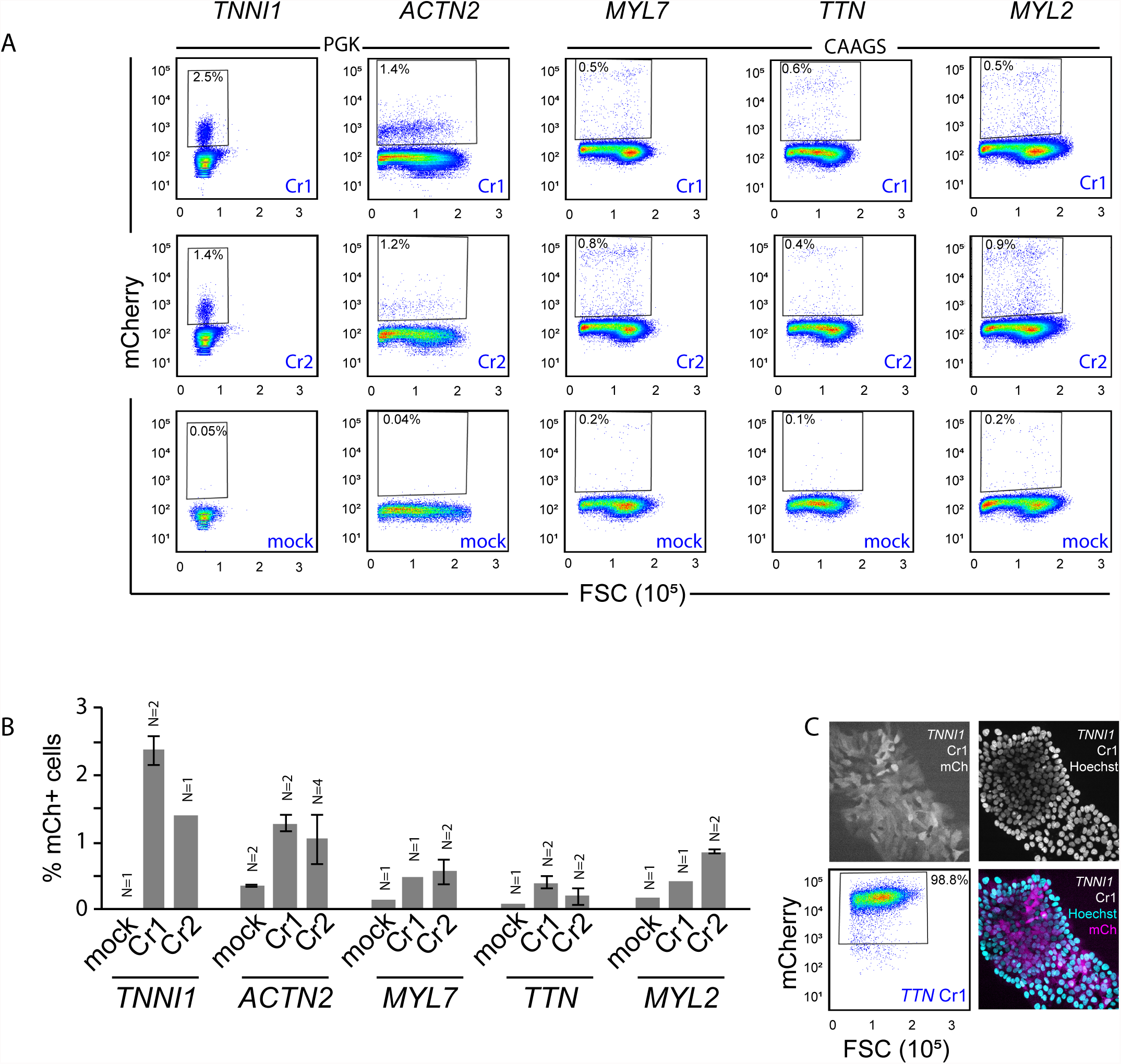
Fluorescence activated cell sorting (FACS) of mCherry-expressing cells to establish the efficacy of multi-step editing at transcriptionally silent loci. **A.** Percentages of mCherry-expressing cells isolated after transfection with donor plasmids in conjunction with two unique crRNAs complexed with Cas9 targeting the intended locus (top and middle row of panels), alongside mock transfections (bottom row). The boxes indicate the gating applied to determine mCherry positive cells used for determining HDR efficiency and sorting. The identity of the targeting crRNA is indicated in blue font within each plot. mCherry fluorescence is indicated on the y-axis and forward scatter (FSC) is indicated on the x-axis. Mock conditions for *TNNI1* and *ACTN2* consisted of the plasmid donor with no crRNA, while for *MYL7*, *TTN* and *MYL2* the mock conditions were with an irrelevant non-targeting crRNA **B.** All data from (A) is displayed in graphical format. Standard deviations are indicated where multiple replicate transfections were performed. **C.** Live imaging and FACS were performed one expansion passage after FACS enrichment in order to validate sorting purity. Example images of mCherry and Hoechst (nuclear dye) from the *TNNI1* Cr1 population after sorting are shown. Cells isolated from the *TTN* Cr1 enrichment experiment were analyzed by flow cytometry and found to be 98.8% pure.

We chose to use two different crRNAs for each target because we previously found that editing success can be highly dependent on the target sequence at a given target locus (Roberts et al. 2017). Both crRNAs for all five genes resulted in successful HDR at a rate (0.4-2.5%) consistent with our previous study (0.2-5%) tagging genes expressed in stem cells. In contrast, experiments performed with mock conditions (non-targeting crRNA, or no crRNA) showed few mCherry-expressing cells (Fig. 2A). We hypothesize that these control conditions reveal the low rate of random, non-targeted integration of the construct into the genome (Fig. 2A, bottom row, B). mCherry-positive cells were sorted and these enriched putatively edited cells were expanded and confirmed to consist of >90% mCherry-positive cells by imaging and/or flow analysis (Fig. 2C for examples, Fig. 3A mock).

**Figure 3.**
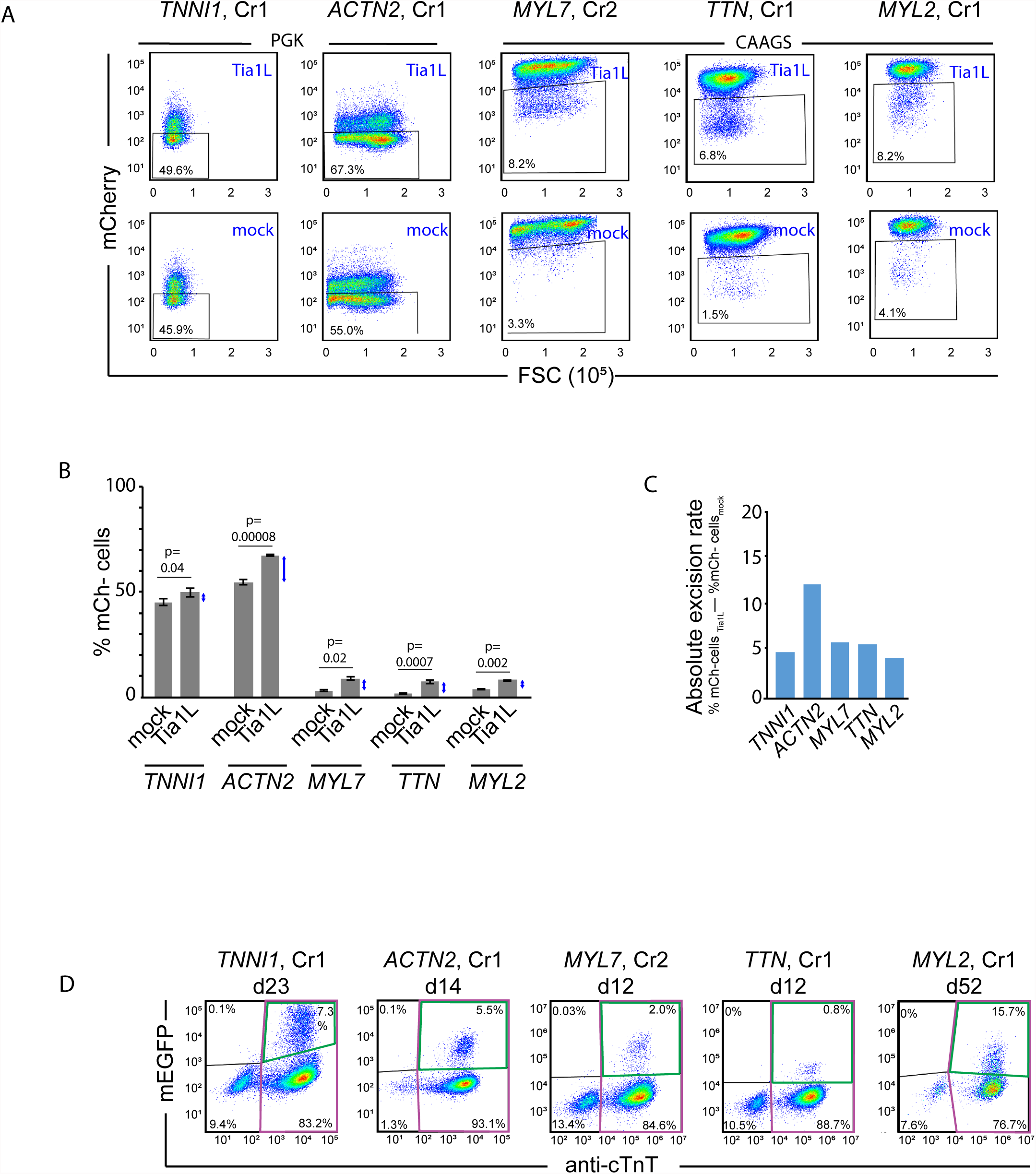
Sorting of mCherry-negative cells to measure excision and obtain putatively mEGFP-tagged cells. **A.** mCherry-expressing cells isolated from targeting experiments (shown in Fig. 2) were transfected after recovery with either Tia1L or mock RNP. mCherry-negative cells were then collected as putatively excised cells from the Tia1L transfected condition. The percentage of mCherry-cells, according to the displayed gates, were measured and are as indicated from both the Tia1L and mock transfected conditions. mCherry fluorescence is indicated on the y-axis and forward scatter (FSC) is indicated on the x-axis. **B**. Percentages of mCherry-negative cells (as determined by FACS threshold shown in panel A) from both mock and Tia1L excision conditions are shown in graphical format. Standard deviation describes variance between replicate conditions. P-values from Student’s t-test are shown, demonstrating significantly elevated abundance of mCherry-negative cells when excision was performed with the Tia1L crRNA. **C.** The percentage of on-target excised cells (blue arrows in panel B) calculated by subtracting the mean percentage of mCherry-negative cells in the Tia1L excised condition from the mean percentage of mCherry-negative cells in the mock excised condition is shown on the y-axis. This value was used as an estimate for absolute rate of excision. **D.** mCherry-negative cells isolated after Tia1L-mediated excision were differentiated into cardiomyocytes and analyzed by flow cytometry for expression of mEGFP and cardiac troponin T (cTnT+, pink box). mEGFP expression was observed within the cardiomyocyte population (cTnT+/mEGFP+, green box). The duration of differentiation before analysis is indicated. The percentage of cTnT+/mEGFP+ (upper right quadrant within each plot) cells is indicated, and was interpreted as a proxy for estimated tagging efficiency.

### Step 2 - Excision of the mCherry selection cassette with CRISPR/Cas9 and NHEJ/MMEJ repair

After FACS enrichment, mCherry-positive, putatively edited cells were subjected to a second round of editing with RNP complexes that targeted the Tia1L target sequences, in order to excise the selection cassette (Fig. 1B, Step 2). Following this transfection, cultures were recovered, expanded over two passages (~7-8 days), and sorted for mCherry-negative cells. The observation of a subset of mCherry-negative cells was indicative of excision, and this population of cells was more prominent in the Tia1L transfected cells compared to the mock crRNA in all experiments (Fig. 3A, B). mCherry excision was performed on populations derived from one targeting crRNA within each experiment.

We initially tested the excision step of this editing strategy with *TNNI1* and *ACTN2*. However, confirming and sorting excised *TNNI1*- and *ACTN2*-targeted cells was challenging due to the low expression of hPGK-driven mCherry in the starting population and the gradual loss of mCherry expression due to silencing prior to excision (Fig. S3). However, the reproducible loss of mCherry expression relative to mock excision controls and subsequent clonal analysis suggested that excision of the selection cassette occurred in a subset of these *TNNI1* and *ACTN2*-edited cell populations (Fig. S3A). In contrast, identifying and sorting the mCherry-negative excised population for *TTN*, *MYL2*, and *MYL7* was easier due to the higher and more stable expression of CAGGS-driven mCherry before excision compared to mCherry driven by hPGK. This is highlighted by the 2-7 fold increase in mCherry negative cells compared to the mock controls for *TTN*, *MYL2*, and *MYL7* (Fig. 3A). We used the differences between the percentages of mCherry-negative cells observed in Tia1L and mock excision conditions to calculate the absolute rate of excision in each of the five experiments, which varied from ~4-12% (Fig. 3C). We also calculated the estimated frequency of alleles in each Tia1L excised population that we hypothesized were candidates for tagging because they represented on-target editing events, rather than random genomic integrations of the donor plasmid. We performed this calculation by dividing the relative difference of mCherry-negative cell abundance by the absolute number of mCherry-negative cells in the Tia1L excised population (Fig. S3B).

### Step 3 - Initial confirmation of mEGFP-tagging within mosaic cardiomyocyte cultures

The mCherry-negative populations were sorted, expanded, and evaluated for expression of the mEGFP-fusion protein upon differentiation into cardiomyocytes (Fig. 1B, Step 3, Fig. 3D). All five gene edited populations resulted in robust cardiomyocyte differentiation with high (>86%) percentages of cells expressing the cardiomyocyte marker cardiac troponin T (cTnT). We identified a subset of mEGFP-expressing cells (1-16%) within the cTnT-positive cardiomyocyte cell populations in all 5 targeting experiments (Fig. 3D). In contrast, mEGFP expression was extremely rare (<0.1%) in cTnT-negative cells, strongly suggesting that mEGFP expression was specific to cardiomyocytes. The expression and sarcomeric localization of the mEGFP fusion protein in these cardiomyocytes was also confirmed by microscopy (data not shown).

### Step 4 - Genetic screening for clones with precisely edited mEGFP-tagged alleles

Encouraged by our ability to identify mEGFP-expressing cardiomyocytes from populations of edited hiPSCs, we set out to identify precisely edited clones (Fig. 1B, Step 4). 150-200 colonies were isolated from each putatively excised population and screened for precise editing similar to previously described methods (Roberts et al. 2017). We performed a sequential genomic screen consisting of a primary multiplexed ddPCR assay to measure the genomic copy number of several key sequences from each clone (mEGFP/AmpR/mCherry) followed by junctional PCR and Sanger sequencing as described below. As excision experiments were only performed on mCherry-expressing cells from one initial transfection, all analyzed clones were derived from edits generated by a single targeting crRNA.

First, we evaluated the presence and genomic copy number of the mEGFP tag sequence in each clone. We interpreted the presence of at least one copy of the mEGFP tag as evidence that the homology donor cassette was stably incorporated into the genome (Fig. 4A, x-axis). The majority of clones contained mEGFP and we selected a subset with one copy, indicative of a monoallelic edit for further analysis. Clones with no mEGFP or mosaic mEGFP (a non-integer genomic copy number) were rejected at this step. No biallelic clones were detected for any of the target genes (Fig. 4A). We also rejected all clones with AmpR and/or mCherry ddPCR signal with the goal of identifying clones that had monoallelic mEGFP, no plasmid backbone incorporation (AmpR-), and excision of the mCherry cassette (mCherry-). In some clones, several genomic copies of mCherry and/or AmpR were observed, although the frequencies of these events varied by gene (Fig. 4A, y-axis). We hypothesized that these ddPCR profiles indicate multiple misediting and sorting outcomes including incorporation of the plasmid backbone sequence into the genome, stable integration at an off-target site during the first editing step, and/or imperfect selection post-excision. Notably, this was rare for *MYL2* where a low rate of attrition was observed at this step (Fig. 4A, B). Using this multiplexed screen, we identified at least two clones from all five targeting experiments with ddPCR signatures indicating the stable presence of a single mEGFP tag copy in the genome, the absence of AmpR (indicator of imprecise HDR) and absence of mCherry (indicator of inefficient excision).

**Figure 4.**
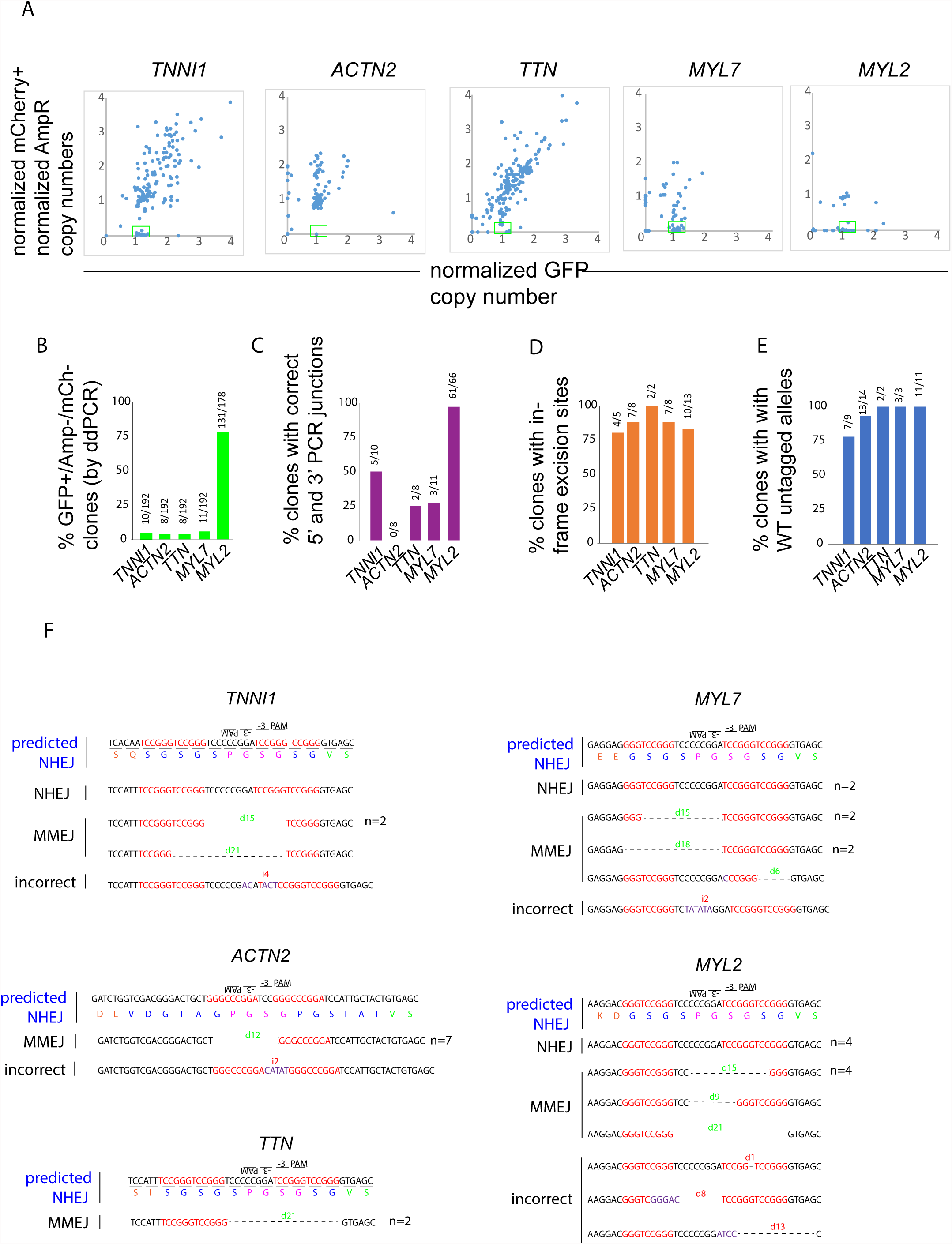
Genetic analysis of precise mEGFP tagging using multi-step targeting and excision in clones. **A.** Clones from each targeting experiment were analyzed according to their normalized genomic copy number of the mEGFP, mCherry and AmpR sequences. Clones were categorized as candidates for further analysis if the mEGFP genomic copy number was consistent with monoallelic or biallelic tagging (copy number of 1 or 2), and additive copy number of AmpR and mCherry was < 0.2. Plots display the normalized mEGFP copy number (x-axis) plotted against the normalized additive mCherry and AmpR copy numbers (y-axis). Clones considered for further analysis are indicated in each plot by the green boxes. **B.** The percentage of clones validated by ddPCR is displayed in bar graphs. **C.** Percentages of ddPCR validated clones with validated PCR junctions (with expected product sizes) between the mEGFP tag and the surrounding genomic region, distal to the homology arms, are shown. **D.** The percentages of analyzed clones with in-frame sequences of peptide linkers at the excision site are additionally shown, demonstrating a high rate of in-frame excision predicted to generate effectively tagged clones. **E.** The percentage of clones with WT untagged alleles is shown, demonstrating the relative low impact of unintended NHEJ at the targeted locus. **F.** In each experiment, peptide linkers at the site of excision were sequenced and are shown. In-frame repair outcomes after excision are indicated. All outcomes within analyzed clones are shown, including aberrant insertions and deletions, and all insertions and deletions are indicated by “i” or “d” followed by the number of nucleotides inserted or deleted, respectively. The frequency of clones for each outcome is indicated to the right of each sequence if greater than one.

We next performed junctional PCR on the mEGFP+/mCherry-/AmpR- clones that were identified as candidates from the ddPCR assay to evaluate editing at the appropriate genomic location. Clones that were rejected because they contained evidence of stable genomic integration of the mCherry and/or AmpR sequences were not further analyzed by junctional PCR or Sanger sequencing in the current report, and we hypothesize that many of these clones were edited at the intended genomic locus but in an incorrect manner. PCR primers were designed to amplify the sequence junctions between the mEGFP tag and the genomic sequences 5’ and 3’ of the homology arms. All mEGFP+/mCherry-/AmpR- clones identified from the ddPCR screen underwent editing at the appropriate genomic locus, as judged by the successful amplification of overlapping PCR products on both sides of the tag sequence insertion (data not shown). The extent to which these PCR products matched the anticipated product size was specific to each experiment (Fig. 4C). Of these, appropriate PCR junction products were observed for 5/10 *TNNI1* clones, 2/8 *TTN* clones, 3/11 *MYL7* clones, and 61/66 *MYL2* clones, and were further considered as candidates (Fig. 4C). All 8 *ACTN2* clones identified by the ddPCR screen contained an aberrantly large 3’ junction due to a duplication in the homology arm region during HDR (data not shown). All clones from all five targeting experiments that displayed aberrantly large PCR junction products were flawed at either the 5’ or the 3’ junction, but not both, supporting the conclusion that HDR misrepair is a strand specific DNA repair event.

Next, Sanger sequencing of the target locus was performed on clones with appropriately sized junctions to test for precise HDR and excision outcomes. The majority of clones confirmed by 5’ junctional PCR contained sequences with predicted NHEJ or MMEJ outcomes following excision (Fig. 4D). In addition to in-frame deletion of the selection cassette, precisely edited clones contained a range of linker-mEGFP fusions made possible by the donor template design. 19% of clones (n=36) contained a Ser-Gly-Pro-Gly (SGPG) linker consistent with an NHEJ outcome (Fig. 4D,F). Other in-frame linkers of differing length arising from 3-21 bp deletions occurred in 64% of clones and were likely a result of MMEJ-driven repair (Fig. 4D,F). We rejected 17% clones due to small out-of-frame deletions in the sequence near the excision site (Fig. 4D,F). Finally, all clones with PCR-validated 3’ junctions (n=23) showed the anticipated sequence identity (data not shown).

In a final assay, we amplified and sequenced the untagged allele in all clones with correct ddPCR signatures and validated junctions, and in some cases, clones with incorrect junctions. NHEJ damage to the untagged allele was observed in 3/39 clones across all five experiments (Fig. 4E), and these clones were rejected. The low rate of observed NHEJ damage to the untagged alleles within HDR-targeted clones was consistent with our previous report (Roberts et al. 2017).

### Step 5 - Confirmation and validation of mEGFP-tagging in hiPSC-derived cardiomyocytes

All clones that satisfied the genomic criteria for precise in-frame mEGFP fusion to the targeted locus after step 4 were subjected to directed cardiomyocyte differentiation to evaluate tagging efficiency of the sarcomeric proteins (Fig. 1B, Step 5). All clones demonstrated robust differentiation into cardiomyocytes with spontaneous contraction and mEGFP expression (Fig. 5A,B, Table 1, Fig. S2). Flow cytometry was performed 12 days after initiating cardiac differentiation and revealed that the majority of cells in all tested *ACTN2, TNNI1*, *TTN*, *MYL7* and *MYL2* clones were cardiomyocytes (>70% cTnT+)(Fig. 5B, 6SB Table 1). This included a single rejected clone from the ACTN2 experiment with low GFP expression in cardiomyocytes (<26% GFP+/cTnT+ cells). With the exception of late-expressing *MYL2*, all other clones expressed mEGFP in the majority of cardiomyocytes (>90% mEGFP+/cTnT+), suggesting that mEGFP-tagged alleles are expressed following cardiomyocyte differentiation (Fig. 5 A,C, Table 1). Consistent with varying transcript abundance with bulk RNA-sequencing (data not shown), the intensity of mEGFP expression was different across the five genes, with the lowest levels of expression observed in *TTN*-mEGFP (Fig. 5A). Expression of *ACTN2*-mEGFP was observed in multiple clones despite imprecise editing (duplication of donor plasmid elements) at the 3’UTR (Fig. 5A-C). Consistent with our previous observation in non-clonal differentiation experiments, cTnT-/mEGFP+ cells within the clonal differentiated cells were extremely scarce (<1.3%) in all clones tested suggesting that expression of the fusion protein was specific to the cardiomyocyte lineage (Fig. 5A).

**Figure 5.**
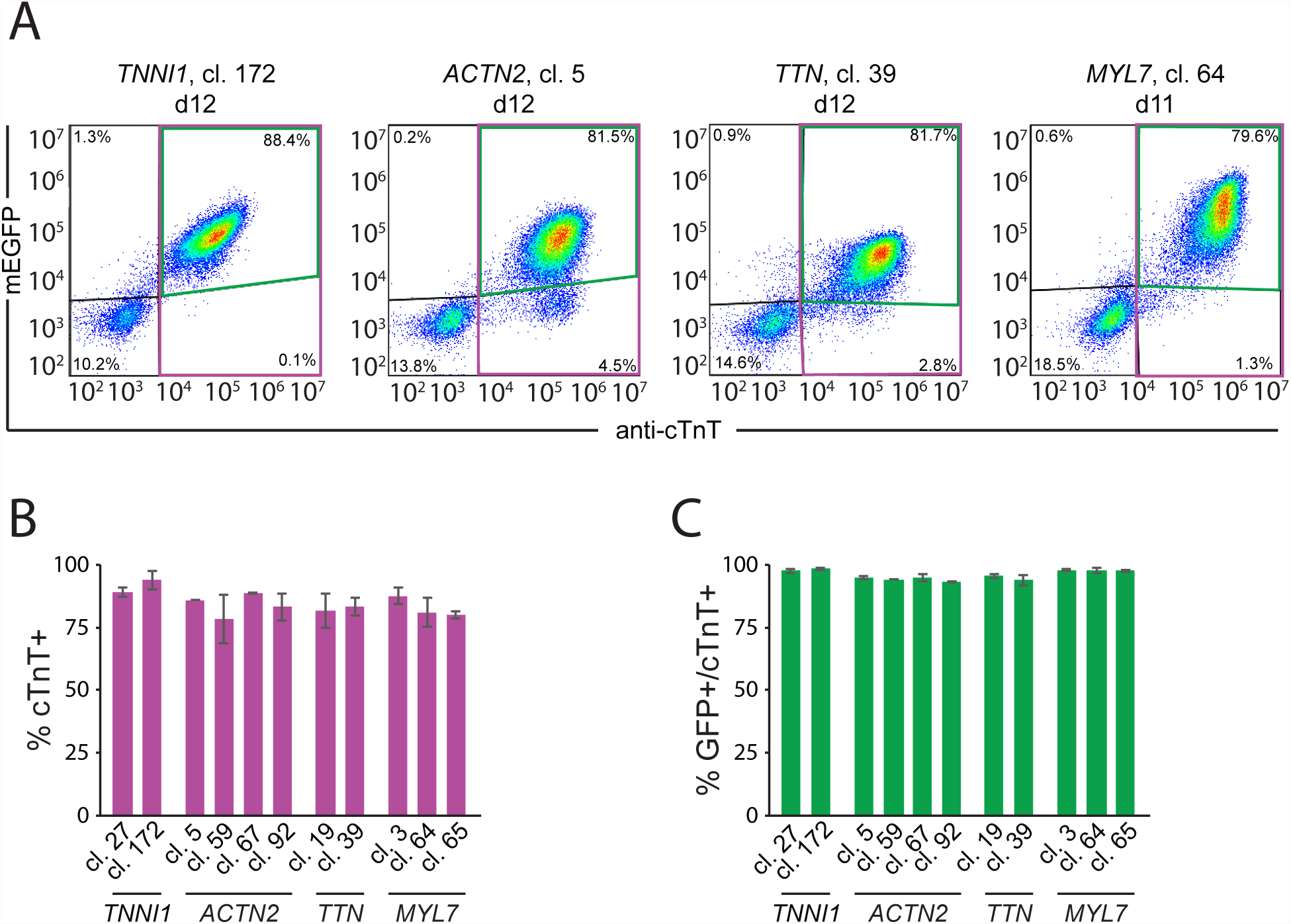
Quantitative assays to evaluate cardiomyocyte differentiation efficiency and mEGFP-tagged protein expression in precisely excised clones. **A.** Validated clones were differentiated into cardiomyocytes. mEGFP fluorescence in fixed cells was measured by flow cytometry (y axis) and plotted against antibody staining intensity for the cardiomyocyte marker cardiac troponin T (cTnT, x axis). Samples were collected for analysis ~12 days after the initiation of differentiation, and plots from representative clones are shown. Each clone displayed successful differentiation, with the majority of cells expressing cTnT (pink box). Within the cTnT+ cell population, a significant percentage of cells expressed mEGFP (cTnT+/mEGFP+, green box). Percentages of cells gated as positive and negative for both mEGFP fluorescence and cTnT staining are as indicated. **B.** The percentages of cells positive for cTnT from precisely edited clones are shown. C. The percentages of cardiomyocytes (cTnT+) expressing mEGFP from edited clones in B are shown. In **B** and **C**, error bars are standard deviation amongst biological replicates. *ACTN2* experiments included two replicates. All other experiments consisted of three replicates.

**Table 1.** Quality control criteria to evaluate the robustness of clonal line differentiation, pluripotency and genomic stability. Clones from each experiment were evaluated for karyotypic abnormalities with metaphase spreads. Flow cytometric analysis of nuclear pluripotency markers is shown, with minimum values obtained among all trials displayed and number of trials in parentheses. Germ layer marker expression was measured by RT-ddPCR for *TNNI1* clone 172 (other clones in progress). Cardiomyocyte differentiation was performed and the percentage of cTnT+ cells, and cTnT+/mEGFP+ cells were measured by flow cytometry. Errors indicate standard deviation. Bold entries are clones identified for public release. *ACTN2* clone 20 (italicized) was rejected due to low mEGFP expression in cTnT+ cardiomyocytes.

|  |  |  |  |  |  |  | Cardiomyocyte differentiation metrics, % positive (n) |  |  |  |  |
| --- | --- | --- | --- | --- | --- | --- | --- | --- | --- | --- | --- |
|  |  |  | Pluripotency markers,<br>% positive (n) |  |  | Germ layer directed<br>differentiation | Cytokine protocol |  |  | Small Molecule protocol |  |
| Cell Line | Clone | Karyotype | Oct3/4 | Nanog | Sox2 | Brachyury, Sox17, Pax6 | cTnT | cTnT/GFP |  | cTnT | cTnT/GFP |
| Unedited |  | Normal | >96 (6) | >99 (6) | >99 (6) | Pass | 91.9 ± 3.9 | na | (5) | 82.7 ± 6.5 | na (10) |
| TNNI1 | 25 | Abnormal, clone discontinued |  |  |  |  |  |  |  |  |  |
|  | 27 | Normal | >96 (2) | >99 (2) | >99 (2) | nd | 89.1 ± 1.9 | 97.7 ± 0.8 | (3) |  |  |
|  | 172 | Normal | >97 (2) | >99 (2) | >99 (2) | Pass | 93.9 ± 3.6 | 98.6 ± 0.4 | (3) |  |  |
| ACTN2 | 5 | Normal | 100 (1) | 100 (1) | 100 (1) | nd | 82.2 ± 12.9 | 93.3 ± 2.7 | (2) | 86.0 ± 0 | 94.8 ± 0.7 (2) |
|  | 20 | Normal | 99 (1) | 100 (1) | 100 (1) | nd | 94.7 ± 0.9 | 25.7 ± 10.5 | (2) |  |  |
|  | 59 | Normal | 100 (1) | 100 (1) | 100 (1) | nd | 84.9 ± 17.4 | 92.9 ± 5.8 | (2) | 78.4 ± 9.8 | 94.1 ± 0 (2) |
|  | 67 | Normal | 98 (1) | 100 (1) | 100 (1) | nd | 81.8 ± 14.3 | 90.6 ± 4.2 | (2) | 88.9 ± 0.1 | 94.9 ± 1.4 (2) |
|  | 92 | Normal | 99 (1) | 100 (1) | 100 (1) | nd | 81.9 ± 12.9 | 90.0 ± 6.2 | (2) | 83.2 ± 5.5 | 93.4 ± 0.1 (2) |
| TTN | 19 | Normal | 97 (1) | 99 (1) | 100 (1) | nd | nd | nd |  | 81.8 ± 6.8 | 95.7 ± 0.8 (3) |
|  | 39 | Normal | 99 (1) | 100 (1) | 100 (1) | nd | nd | nd |  | 83.3 ± 3.4 | 94.0 ± 2.1 (3) |
| MYL7 | 3 | Normal | 100 (1) | 100 (1) | 100 (1) | nd | nd | nd |  | 87.5 ± 3.3 | 97.9 ± 0.4 (3) |
|  | 64 | Normal | 100 (1) | 100 (1) | 100 (1) | nd | nd | nd |  | 81.0 ± 5.7 | 97.8 ± 1.2 (3) |
|  | 65 | Normal | 100 (1) | 100 (1) | 100 (1) | nd | nd | nd |  | 80.0 ± 1.3 | 97.6 ± 0.5 (3) |
| MYL2 | 27 | Normal | 100 (1) | 100 (1) | 100 (1) | nd | nd | nd |  | 87.6 ± 3.3 | 5.0 ± 3.6 (2) |
|  | 67 | Normal | 100 (1) | 100 (1) | 100 (1) | nd | nd | nd |  | 70.2 ± 21.9 | 2.4 ± 0.2 (2) |
|  | 68 | Normal | 99 (1) | 100 (1) | 100 (1) | nd | nd | nd |  | 84.3 ± 1.6 | 3.0 ± 3.6 (2) |
|  | 70 | Normal | 99 (1) | 100 (1) | 100 (1) | nd | nd | nd |  | 90.7 ± 1.2 | 0.9 ± 0.1 (2) |
|  |  |  |  |  |  |  | Samples collected at days 14-21 |  |  | Samples collected at days 12-15 |  |

Samples collected at days 14-21
Samples collected at days 12-15

### Step 6 - Imaging of clonal mEGFP-tagged hiPSC-derived cardiomyocytes

In addition to confirming expression of mEGFP by flow cytometry, cardiomyocytes generated from all five gene edited clonal lines were re-plated on glass-bottom, PEI/laminin-coated plates to perform live-cell imaging and immunocytochemistry (Fig. 1B, Step 6, Fig. 6). The five cell lines each represent different structural elements of the cardiomyocyte sarcomere and thus all fusion proteins they encode should localize to the sarcomere but with different localization patterns, depending on their role. Live imaging revealed sarcomeric localization for all of the mEGFP-tagged fusion proteins, each one displaying a striated appearance (Fig. 6A). However, the exact nature of the striations differed as expected for each of these sarcomeric proteins. The thick and thin filament fusion proteins from the *MYL7* and *TNNI1* experiments localize to a thicker banding pattern of m EGFP signal interspersed with thinner “dark” lines where mEGFP is absent. This is consistent with their localization between z-discs in the myofibril (Luther 2009). In contrast fusion proteins from the *ACTN2* and *TTN* experiments both localized to a thinner banding pattern within the sarcomere, representing the Z-disc and the M-line respectively. The banding pattern for *TTN* was especially revealing as this protein spans the entire length of the sarcomere but is only mEGFP tagged at the C-terminus, which is localized at the M-line.

**Figure 6.**
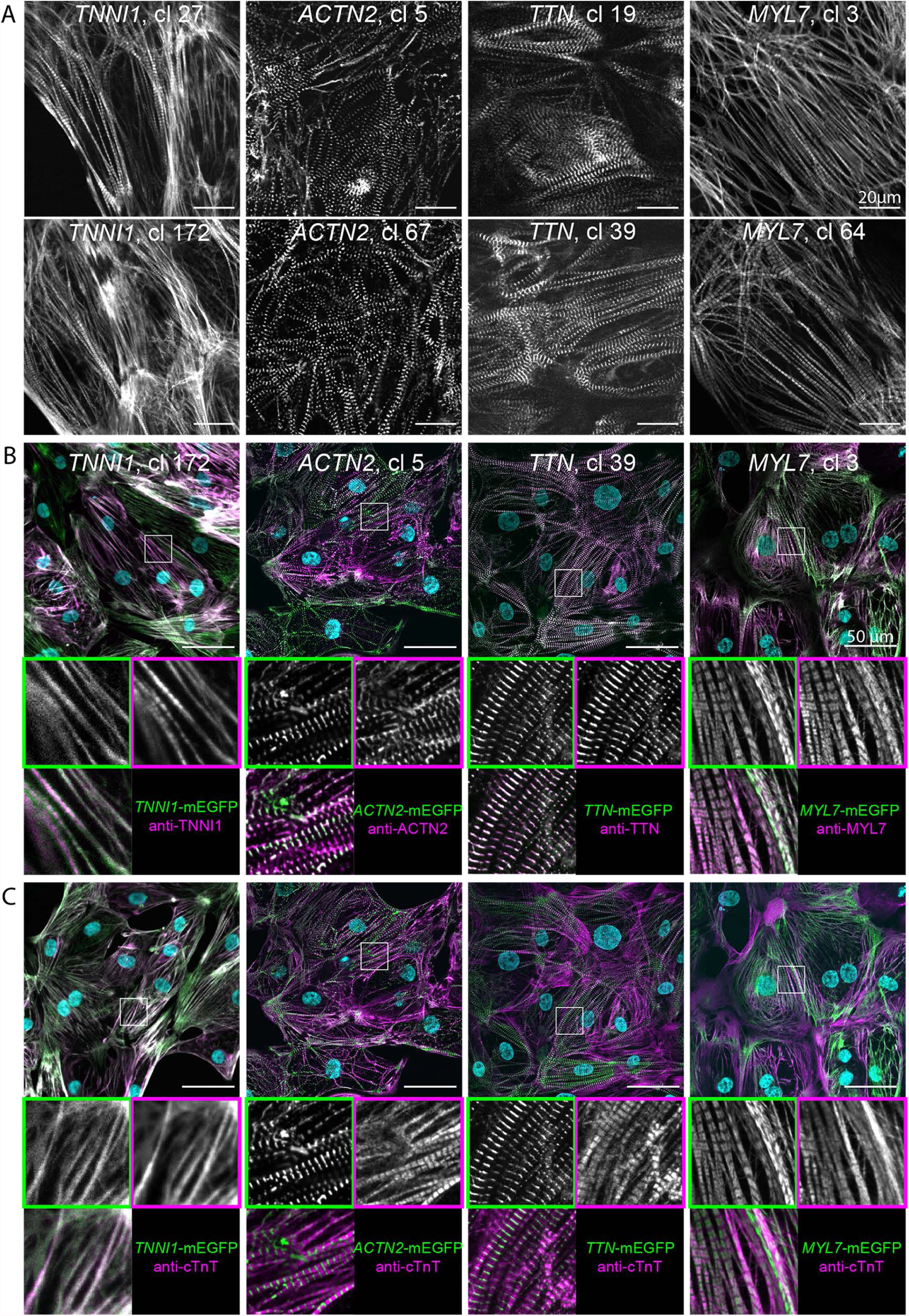
Imaging experiments to evaluate sarcomeric localization of the mEGFP-tagged proteins. **A.** Cardiomyocytes from two independently edited clones for each of the five targeting experiments were plated on glass bottomed plates coated with PEI/laminin and were imaged live using spinning disc confocal microscopy for mEGFP fluorescence. Scale bars: 20 µm. **B.** Representative images of mEGFP fluorescence and antibody labeling for the protein product of the indicated gene are shown for a representative clone for each target. For all multi-channel images, mEGFP signal is shown in green, antibody label is shown in magenta and DAPI stain (DNA) is shown in cyan. Scale bars: 50 µm. Insets are 25 µm and show mEGFP signal (upper left, green box), antibody labeling signal (upper right, magenta box) and merge (bottom left). **C.** Expression of mEGFP is observed specifically in cardiomyocytes. Representative images of mEGFP signal and the cardiomyocyte marker cTnT indicate that mEGFP expression is observed in cTnT-positive cells. Scale bars: 50 µm. Insets are 25 µm and show mEGFP (upper left, green box), anti-cTnT (upper right, magenta box) and merge (bottom left). All cardiomyocytes shown here were imaged approximately 20 days after initiating differentiation.

Antibodies specific to the targeted proteins co-localized with mEGFP, confirming appropriate localization of the tagged protein in cardiomyocytes (Fig. 6B). Unedited cells were also fixed and immunolabeled with antibodies specific to each target, and labeling patterns were found to be comparable to that in the mEGFP-tagged cells (data not shown). We also confirmed expression of cTnT in edited cardiomyocytes (Fig. 6C). The vast majority of cTnT-positive cells were also mEGFP+, even after 3-4 weeks of differentiation, indicating that expression of the mEGFP-tagged allele in these cardiomyocytes remained stable and clonal over time.

*MYL2*-mEGFP tagged clones were not evaluated according to the same criteria in the current report due to their requirement for evaluation at a later time point following differentiation (pending studies). However, we observed that up to 33% of the cTnT-positive cells were GFP-positive after 26 days of differentiation, with some variance between analyzed clones (Fig. S6A-C). Where observed, GFP signal within these clones localized between the z-discs in the myofibril, as expected (Fig. S6D).

## Discussion

Endogenous fluorescent tagging in hiPSCs enables live imaging to study the organization and dynamics of key proteins and structures in stem cells and their derivatives. The advent of efficient and accessible gene editing tools like CRISPR/Cas9 has only recently merged endogenous tagging approaches with the differentiation potential of hiPSCs, as demonstrated in a recent study evaluating adhesion in sarcomere assembly using paxillinm-EGFP tagged cardiomyocytes (Chopra et al. 2018). This more recent approach using pluripotent models follows established *in vivo* approaches with knock-in mice expressing endogenously tagged *MYL2* and *TTN* that have revealed basic mechanisms of sarcomere dynamics (da Silva Lopes et al. 2011, Ishizu et al. 2017). Partnering insights gained from *in vivo* studies with hiPSC-derived models will reveal mechanisms important for differentiation, development and disease. However, such approaches require systematic and scalable strategies for endogenous tagging, irrespective of transcriptional activity of the target gene in hiPSCs.

To do this, we developed a unique multi-step CRISPR/Cas9-mediated editing strategy for tagging non-expressed genes in hiPSCs and used this methodology to tag several genes expressed specifically during cardiomyocyte differentiation. Our donor plasmid design enabled us to detect HDR at targeted non-expressed loci, enrich for putatively tagged cells with a constitutively expressed selection cassette, and generate mEGFP fusion alleles lacking genomic scars using a Cas9/MMEJ-driven excision strategy. This approach uniquely relies on CRISPR/Cas9 for both the incorporation and subsequent excision of the selection cassette and provides the added benefits of selection without drugs. Our ability to confirm delivery of the selection cassette via HDR, enrich for mCherry expression, excise the selection cassette, and incorporate a tunable linker in multiple clones derived from all five edited hiPSC lines demonstrates the feasibility of this method and workflow.

We previously described mEGFP-tagging experiments targeting 10 expressed genes in WTC hiPSCs using a similar HDR delivery protocol, but with donor plasmids designed to deliver only an in-frame mEGFP tag at the N- or C-terminus of the target gene. HDR rates typically ranged from 0.2-5% for these transcriptionally active loci in hiPSCs. We observed similar rates (0.4-2.5%) in the current study using a much larger donor plasmid, suggesting that targeting non-expressed rather than expressed loci with a selection cassette much larger than the tag alone permits similar HDR rates for silent editing. Testing multiple crRNAs in parallel for each target also ensured editing success. In both studies, >90% edited cells were recovered by FACS, highlighting the utility of a drug-free enrichment strategy, especially when HDR rates are low. CRISPR/Cas9 was utilized a second time to excise the mCherry selection cassette from the enriched population of edited cells. The relative efficiency of this step (~4-12%) varied by gene and promoter with the CAGGS promoter (*TTN*, *MYL2* and *MYL7*) preferable to hPGK (*ACTN2*, *TNNI1*). In all cases, the rate of excision was sufficient for robust FACS enrichment and exceeded the rate of HDR observed in the initial delivery step.

Successful gene tagging was achieved with this methodology for all five target genes with a significant number of clones containing mEGFP at the intended locus. Further screening by PCR and Sanger sequencing identified a subset of clones with precise editing as defined by the absence of any mutations or duplications at either the tagged or untagged allele. The frequency of precise editing, as determined by the percentage of clones validated in our ddPCR screen, varied by target locus, much like in our previously reported data set targeting expressed loci (Roberts and Haupt et al. 2017). Since the same Tia1L sequences were used for excising the selection cassette in the second step of this method in all five experiments, we hypothesize that the variability in editing precision is introduced during the first editing event of HDR, which is influenced by the target locus and crRNA. We plan to further investigate the association between HDR and precision and increase the frequency of generating precisely edited and excised clonal lines.

We also demonstrate the use of MMEJ sequences to guide scarless excision with a tunable linker as well as introduction of a canonical peptide linker for when DNA repair is mediated by NHEJ in multiple clones. These features provide flexibility for adding a specific linker based on known properties of the target protein and provides a strategy to precisely engineer various edited outcomes. This can include the addition of an epitope tag, a cleavable peptide, or no intervening sequence as demonstrated for correcting disease mutations in a recent study (Kim et al. 2018). Our results with these five target genes suggest that the use of a CAGGS-driven selection cassette along with specific microhomology sequences provide means of introducing scarless edits at silent loci. To our knowledge, this represents the first report utilizing MMEJ with CRISPR/Cas9-mediated endogenous tagging to generate scarless edits and custom linkers at silent loci.

All precisely edited clonal cell lines differentiated into cardiomyocytes with timely expression of the mEGFP-fusion protein, demonstrating endogenous regulation of the tagged protein during cardiac differentiation. Appropriate expression and localization was observed in tagged *ACTN2* clones despite imprecise editing at the 3’UTR, suggesting that the compromised UTR does not affect the expression or localization of *ACTN2*-mEGFP. Tracking the expression and localization of the five sarcomeric proteins tagged in this report will help evaluate the expression and dynamics of each fusion protein during the differentiation and maturation process, as well as validate the integrity of the tagged alleles. Since the five tagged genes encode important sarcomeric proteins, successful tagging will enable live cell analysis of structural organization during differentiation and development.

Here we have demonstrated the feasibility of a multistep editing procedure to tag silent genes in hiPSC in a scarless manner. The ability to generate multiple mEGFP-tagged clonal hiPSC lines for cardiomyocyte-specific genes shows the utility and flexibility of this approach. This strategy may be adapted for editing genes associated with development and disease in various cell types differentiated from hiPSCs.

## Materials and Methods

### Cell Culture

All work with human induced pluripotent stem cell (hiPSC) lines was approved by internal oversight committees and performed in accordance with applicable NIH, NAS, and ISSCR Guidelines. The WTC hiPSC cell line was generated by Dr. Bruce Conklin (The Gladstone Institutes) and maintained using described methods (Kreitzer et al. 2013). Edited cell lines described in this report can be obtained by visiting the Allen Cell Explorer (Allen Institute for Cell Science 2017). A detailed protocol outlining this method can be found at allencell.org.

### Donor Plasmids, crRNAs and Cas9 Protein

Donor plasmids were designed uniquely for each target locus. Homology arms 5’ and 3’ of the desired insertion site were each 1 kb in length and designed using the GRCh38 reference genome. WTC-specific variants (SNPs and INDELs) were identified from publicly available exome data (UCSC Genome Browser) and also internal exome and whole genome sequencing data. In cases where the WTC-specific variant was heterozygous, the reference genome variant was used in the donor plasmid; when the WTC-specific variant was homozygous, the WTC-specific variant was used in the donor plasmid. Linkers for each protein were unique to each target and were included in the donor plasmid with microhomology-containing redundancies, such that after MMEJ, the restored sequence would function as a linker 5’ of mEGFP in each C-terminal tagging experiment. The linker sequence was designed to flank Tia1L crRNA binding sites, which in turn flanked the mCherry expression cassette sequence containing either the PGK promoter or the CAGGS. To prevent crRNAs from targeting the donor plasmid sequence, mutations were introduced to disrupt Cas9 recognition or crRNA binding; when possible, these changes did not affect the amino acid sequence. In the ideal case when mutations were unnecessary due to the protospacer spanning the intended genomic insertion site, no mutations were made in the homology arms of the donor plasmid. The plasmids were synthesized and cloned into a pUC57 backbone and prepared for transfection by Genewiz. Custom synthetic crRNAs and their corresponding tracrRNAs were ordered from either IDT or Dharmacon. Recombinant wild type *Streptococcus pyogenes* Cas9 protein was purchased from the UC Berkeley QB3 Macrolab. All targeting experiments discussed here used the mEGFP (K206A) sequence. Detailed information relating to editing design can be found at The Allen Cell Explorer (Allen Institute for Cell Science 2017).

### Transfection and Enrichment by Fluorescence-Activated Cell Sorting (FACS)

Cells were dissociated into single-cell suspension using Accutase as previously described (Roberts and Haupt et al. 2017). Transfections were performed using the Neon transfection system (ThermoFisher Scientific). A detailed protocol for the RNP transfection can be also be found at the Allen Cell Explorer (Allen Institute for Cell Science 2017). Briefly, 8×10^5^ cells were resuspended in 100 µL Neon Buffer R with 2 µg donor plasmid, 2 µg Cas9 protein, and duplexed crRNA:tracrRNA in a 1:1 molar ratio to Cas9. Prior to addition to the cell suspension, the Cas9/crRNA:tracrRNA RNP was precomplexed for a minimum of 10 min at room temperature. Electroporation was with one pulse at 1300 V for 30 ms. Cells were then immediately plated onto growth factor reduced (GFR) Matrigel-coated 6-well dishes with mTeSR1 media supplemented with 1 % Penicillin/Streptomycin (P/S) and 10 µM ROCK inhibitor. Transfected cells were cultured as previously described for 3-4 days until the transfected culture had recovered to ~70% confluence. Negative control transfections were performed in all experiments with the crRNA targeting the AAVS1 locus (protospacer + PAM: 5’ ggggccactagggacaggatTGG 3’) to assess the relative rate of random donor cassette incorporation. Cells were cultured for two passages over 7-9 days before analysis, in order to allow mCherry expression from the episomal donor plasmid to decline. Cells were harvested for FACS using Accutase as previously described. The cell suspension (0.5 – 1.0×10^6^ cells/mL in mTeSR1 with ROCK inhibitor) was filtered through a 35 µm mesh filter into polystyrene round bottomed tubes. Cells were sorted using a FACSAriaIII Fusion (BD Biosciences) with a 130 µm nozzle and FACSDiva software (BD Biosciences). Forward scatter and side scatter (height versus width) were used to exclude doublets and the mCherry+ gate was set using live, negative control transfected cells. Sorted populations were plated into GFR Matrigel-coated 96-well plates (<2×10^3^ cells recovered) or 24-well plates (<1×10^4^ cells recovered) for expansion. The expanded mCherry+ sorted populations were dissociated into single-cell suspension and transfected as previously described with Tia1L-targeted RNP complexes to excise the selectable cassette. Transfected cells were expanded over two passages and sorted as previously described, except that a mCherry-negative sort gate was set using live, mock-excised cells. Sorted populations were expanded in GFR Matrigel-coated tissue culture plates for clonal cell line generation. The Tia1L protospacer + PAM was: 5’ ggtatgtcggg aacctctccGGG 3’.

### Clonal Cell Line Generation

FACS-enriched populations of edited cells were seeded at a density of 1×10^4^ cells in a 10 cm GFR Matrigel-coated tissue culture plate. After 5-7 days clones were manually picked with a pipette and transferred into individual wells of 96-well GFR Matrigel-coated tissue culture plates with mTeSR1 supplemented with 1% P/S and 10 µM ROCK inhibitor for 1 day. After 3-4 days of normal maintenance with mTeSR1 supplemented with 1% P/S, colonies were dissociated with Accutase and transferred into a fresh GFR Matrigel-coated 96-well plate. After recovery, the plate was passaged into daughter plates for ongoing culture, freezing, and gDNA isolation. To cryopreserve clones in a 96-well format, cells were dissociated once they reached 60-85% confluency and pelleted in 96-well V-bottom plates. Cells were then resuspended in 60 µL mTeSR1 supplemented with 1% P/S and 10 µM ROCK inhibitor. Two sister plates were frozen using 30 µL cell suspension per plate, added to 170 µL CryoStor^®^ CS10 (Sigma) in untreated 96-well tissue culture plates. Plates were sealed with Parafilm and stored at −80°C for up to three months.

### Genetic Screening with Droplet Digital PCR (ddPCR)

During clone expansion, >1500 cells were pelleted from a 96-well plate for total gDNA extraction using the PureLink Pro 96 Genomic DNA Purification Kit (ThermoFisher Scientific). ddPCR was performed using the Bio-Rad QX200 Droplet Reader, Droplet Generator, and QuantaSoft software. The reference assay for the 2-copy, autosomal gene RPP30 was purchased from Bio-Rad (assay ID dHsaCP1000485, cat. # 10031243). For primary ddPCR screening the assay consisted of three hydrolysis probe-based PCR amplifications targeted to three different genes: GFP (insert), AMP or KAN (backbone), and the genomic reference RPP30. The following primers were used for the detection of GFP (5’-GCCGACAAGCAGAAGAACG-3’, 5’-GGGTGTTCTGCTGGTAGTGG-3’) and hydrolysis probe (/56-FAM/AGATCCGCC/ZEN/ACAACATCGAGG/3IABkFQ/). This assay was run in duplex with the genomic reference RPP30-HEX. The PCR for detection of the AMP gene used the primers (5’- TTTCCGTGTCGCCCTTATTCC −3’, 5’- ATGTAACCCACTCGTGCACCC −3’) and hydrolysis probe (/5HEX/TGGGTGAGC/ZEN/AAAAACAGGAAGGC/3IABkFQ/). The PCR for detection of the mCherry gene used the primers (5’-TGGCCATCATCAAGGAGTTC −3’, 5’-CTTGGTCACCTTCAGCTTGG-3’) and hydrolysis probe (/5FAM/TCAAGGTGC /ZEN/ACATGGAGGGC/3IABkFQ/). PCR reactions were prepared using the required 2x Supermix for probes with no UTP (Bio-Rad) with a final concentration of 400 nM for primers and 200 nM for probes, together with 10 units of HindIII and 3 µL of sample (30-90 ng DNA) to a final volume of 25 µL. Each reaction prior to cycling was loaded into a sample well of an 8-well disposable droplet generation cartridge followed by 70 µL of droplet generator oil into the oil well (Bio-Rad). Droplets were then generated using the QX200 droplet generator. The resulting emulsions were then transferred to a 96-well plate, sealed with a pierceable foil seal (Bio-Rad), and run to completion on a Bio-Rad C1000 Touch thermocycler with a Deep Well cycling block. The cycling conditions were: 98°C for 10 m in, followed by 40 cycles (98°C for 30 s, 60°C for 20 s, 72°C for 15 s) with a final inactivation at 98°C for 10 min. After PCR, droplets were analyzed on the QX200 and data analysis was preformed using QuantaSoft software. The AMP or KAN signal was determined to be from residual non-integrated/background plasmid when the ratio of AMP/RPP30 or KAN/RPP30 fell below 0.2 copies/genome, because this was the maximum value of non-integrated plasmid observed at the time point used for screening in control experiments (data not shown). For primary screening the ratios of (copies/µL_mEGFP_)/(copies/µL_RPP30_) were plotted against [(copies/µL_AMP_)/(copies/µL_RPP30_) + [(copies/µL_mCherry_)/(copies/µL_RPP30_) to identify cohorts of clones for ongoing analysis. A detailed protocol for these methods is available at the Allen Cell Explorer (Allen Institute for Cell Science 2017).

### Genetic Screening with Tiled Junctional PCR and for Wild Type Untagged Allele Sequences

We evaluated clones with PCR using primers spanning the junction between the tag sequence and the endogenous genomic sequence distal to the homology arm in all GFP+/mCh-/AmpR- clones from each targeting experiment. PCR was used to amplify the tagged allele in two tiled reactions spanning the left and right homology arms, the mEGFP and linker sequence, and portions of the distal g enom ic region 5’ of the left homology arm and 3’ of the right homology arm using gene-specific primers. Both tiled junctional PCR products were Sanger sequenced (Genewiz) bidirectionally with PCR primers when their size was validated by gel electrophoresis and/or fragment analysis (Advanced Analytics Technologies, Inc. Fragment Analyzer). PCR was also used to amplify the untagged allele using gene-specific primers. These primers did not selectively amplify the unmodified locus, but rather amplified both untagged and tagged alleles. Tracking of insertions and deletions (INDELs) by decomposition (TIDE) analysis was performed manually on the amplification reaction after bidirectional Sanger sequencing in order to determine the sequence of the untagged allele. All PCR reactions were prepared using PrimeStar^®^ (Takara) 2x GC buffer, 200 µM DNTPs, 1 unit PrimeStar^®^ HS polymerase, 800 nM primers, 10 ng gDNA in a final volume of 25 µL. Cycling conditions were as follows (98°C 10 s, 70°C 5 s, 72°C 60 s) x 6 cycles at −2°C/cycle annealing temperature, (98°C 10 s, 54°C 5 s, 72°C 60 s) x 32 cycles, 12°C hold. Gene specific primers are included in table.

### *In Vitro* Directed Differentiation of hiPSCs to Cardiomyocytes

Cardiomyocyte differentiation was achieved using a small molecule differentiation protocol similar to previously reported methods, with optimizations to small molecule concentration and timing (Lian et al. 2013). TNNI1 populations and clones and ACTN2 populations were differentiated using previously described methods (Roberts et al. 2017) using a cytokine protocol (Palpant et al. 2015). A detailed protocol can be found at the Allen Cell Explorer (Allen Institute for Cell Science). Briefly, cells were seeded onto GFR Matrigel-coated 6-well tissue culture plates at a density ranging from 0.15-0.25×10^6^ cells per well in mTeSR1 supplemented with 1% P/S and 10 µM ROCK inhibitor, designated as day −3. Cells were grown for three days, with daily mTeSR1 media changes (day −2 and day −1). The following day (designated as day 0), directed cardiac differentiation was initiated by treating the cultures with 7.5 µM CHIR99021 (Cayman Chemical) in RPMI media (Invitrogen) containing insulin-free B27 supplement (Invitrogen). After 48 hours (day 2), cultures were treated with 7.5 µM IWP2 (R&D systems) in RPMI media containing insulin-free B27 supplement. On day 4, cultures were treated with RPMI media containing insulin-free B27 supplement. From day 6 onwards, media was replaced with RPMI media supplemented containing B27 containing insulin (Invitrogen) every 2-3 days.

### Preparation of Glass Bottomed Plates for Cardiomyocyte Re-plating

To prepare 24-well glass bottomed plates for cardiomyocyte plating, plates were treated with 0.5M glacial acetic acid (Fisher Scientific) at room temperature for 20-60 minutes and washed three times with sterile milliQ (MQ) water. Wells were treated with 0.1% PEI (Sigma Aldrich) solution in sterile MQ water overnight at 4°C, then rinsed 2 times with DPBS and one time with sterile MQ water. Finally, wells were incubated overnight at 4°C with 25 µg/mL natural mouse laminin (Thermo Fisher Scientific) diluted in sterile MQ water. Laminin solution was removed immediately prior to re-plating.

### Cardiomyocyte Harvesting, Re-plating and Flow Cytometry Analysis

Cells were harvested using TrypLE Select (10x, Invitrogen) diluted to 2x with Versene (Invitrogen), pre-warmed to 37°C. Cells were washed twice with PBS and incubated with 2x TrypLE/Versene for 6-10 min at 37°C. Cells were gently triturated 8-12x, collected into a tube containing RPMI media (Invitrogen) containing B27 with insulin (Invitrogen), 10 µM ROCK inhibitor, and 200 U/mL DNase I (Millipore Sigma) and pelleted at 211g for 3 min at room temperature. Cells were resuspended in the same media and a 10 µL aliquot was used to count cardiomyocytes in a hemocytometer (INCYTO C-Chip™). Cells were seeded onto PEI/Laminin coated 24-well glass bottomed plates at a density ranging from 0.35-0.75×10^5^ cells per well in RMPI media containing B27 with insulin and 10 µM ROCK inhibitor. 24 hours after plating, media was changed to RPMI media containing B27 with insulin. Imaging was performed 5-21 days after plating, as indicated in figure legends.

To measure cardiac Troponin T (cTnT) expression, cells remaining after plating were pelleted at 211g for 3 min at room temperature. Cells were fixed with 4% paraformaldehyde in DPBS for 10 min at room temperature, washed once with DPBS and then resuspended in 5% FBS in DPBS and stored at 4°C until staining. A maximum of 5 days elapsed between fixation and staining. Fixed cells were incubated in 1x BD P erm/Wash™ buffer containing anti-cTnT AlexaFluor^®^ 647 or equal mass of mIgG1, k AF647 isotype control (all BD Biosciences) for 30 min at room temperature (see antibody table). After staining, cells were washed with BD P erm /Wash™ buffer, then 5% FBS in DPBS and resuspended in 5% FBS in DPBS with DAPI (2µg/mL). Cells were acquired on a CytoFLEX S (Beckman Coulter) or FACSAriaIII Fusion (BD Biosciences) and analyzed using FlowJo software V.10.2. (Treestar, Inc.). Nucleated particles were identified as a sharp, condensed peak on a DAPI histogram and were then gated to exclude doublets using FSC-H/W and, where applicable, SSC-H/W. The cTnT+ positive gate was set to include 1% of cells in the isotype control sample.

### Cardiomyocyte Immunolabeling for Imaging

Fixed cell imaging was performed on cells fixed with 4% paraformaldehyde in PBS with 0.4% Triton X-100 for 10 min at room temperature. Cells were blocked with 1.5% normal goat serum (NGS) with 0.4% Triton X-100 in PBS (blocking solution) for 1-3 hours at room temperature. Primary antibodies were diluted in blocking solution and incubated at 4°C overnight. Secondary antibodies were diluted in blocking solution and applied at 1:500-1:1000 for 2-3 hours at room temperature. Finally, a DAPI counterstain (NucBlue Fixed Cell ReadyProbes Reagent ThermoFisher R37606) was applied for 5 min at room temperature.

### Imaging of Live and Fixed Cardiomyocytes

Live cell imaging was performed on a Zeiss spinning-disk microscope with a Zeiss 20x/0.8 NA Plan-Apochromat, or 40x/1.2 NA W C-Apochromat Korr UV Vis IR objective, a CSU-X1 Yokogawa spinning-disk head, and Hamamatsu Orca Flash 4.0 camera. Fixed cell imaging was done on a 3i spinning-disk microscope with a Zeiss 20x/0.8 NA Plan-Apochromat, or 63x/1.2 NA W C-Apochromat Korr UV Vis IR objective, a CSU-W1 Yokogawa spinning-disk head, and Hamamatsu Orca Flash 4.0 camera. Microscopes were outfitted with a humidified environmental chamber to maintain cells at 37°C with 5% CO_2_ during live imaging.

### G-banding Karyotype Analysis

Karyotype analysis was performed by Diagnostic Cytogenetics Inc. (DCI). A minimum of 20 metaphase cells per clone were analyzed for both ploidy and structural abnormalities.

### Testing Trilineage Potential

To determine differentiation potential, each cell line was differentiated using the STEMdiff™ Trilineage Differentiation kit (STEMCELL Technologies). Cells were plated and differentiated according to the manufacturer’s protocol. Total RNA was harvested on the day of plating and at the final time points specified by the manufacturer, day 5 for Mesoderm/Endoderm and day 7 for Ectoderm. cDNA synthesis was performed with the iScript kit (BioRad). cDNA was assayed by ddPCR for the abundance of germ layer specific markers Brachyury (Mesoderm), Pax6 (Ectoderm), and Sox17 (Endoderm). Transcript levels were normalized to HPRT1 and qualitatively compared to the transcript levels of the AICS-0 parental line. ddPCR primers and probes are included in primer table.

### Expression of Stem Markers by Flow Cytometry

Cells were dissociated with Accutase as previously described, fixed with CytoFix Fixation Buffer™ (BD Biosciences) for 30 min at 4°C, and frozen at −80°C in KnockOut™ Serum Replacement (Gibco) with 10% DMSO. After thawing, cells were washed with 2% FBS in DPBS and permeabilized with 0.1 % Triton X-100 and 2% FBS in DPBS for 20 min at room temperature followed by staining with anti-Nanog AlexaFluor^®^ 647, anti-Sox2 V450, and anti-Oct3/4 Brilliant Violet™ 510 (all BD Biosciences) for 30 min at room temperature. Cells were washed with BD Perm /Wash™ buffer, then 2% FBS in DPBS and before being resuspended in 2% FBS in DPBS for acquisition. Cells were acquired on a FACSAriaI I I Fusion (BD Biosciences) equipped with 405, 488, 561, and 637 nm lasers and analyzed using FlowJo V.10.2 (Treestar, Inc.). Cells were distinguished from debris using forward scatter area vs. side scatter area. Doublets were excluded using forward scatter and side scatter (height vs. width). Positive staining thresholds were established using fluorescence minus one (FMO) controls in which all staining reagents are included except the reagent of interest. For each reagent of interest, the positive gate was set to include 1 % of FMO control cells. The cells stained with the reagent of interest that fell within this gate were used to calculate the number of positive cells. Antibodies used are in table below.

### Antibodies Used for All Studies

**Primary Antibodies**

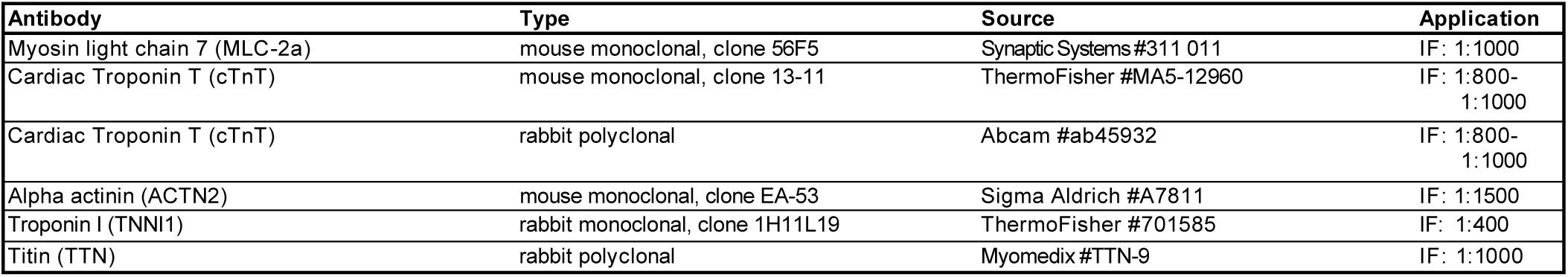

| Antibody | Type | Source | Application |
| --- | --- | --- | --- |
| Myosin light chain 7 (MLC-2a) | mouse monoclonal, clone 56F5 | Synaptic Systems #311 011 | IF: 1:1000 |
| Cardiac Troponin T (cTnT) | mouse monoclonal, clone 13-11 | ThermoFisher #MA5-12960 | IF: 1:800-1:1000 |
| Cardiac Troponin T (cTnT) | rabbit polyclonal | Abcam #ab45932 | IF: 1:800-1:1000 |
| Alpha actinin (ACTN2) | mouse monoclonal, clone EA-53 | Sigma Aldrich #A7811 | IF: 1:1500 |
| Troponin I (TNNI1) | rabbit monoclonal, clone 1H11L19 | ThermoFisher #701585 | IF: 1:400 |
| Titin (TTN) | rabbit polyclonal | Myomedix #TTN-9 | IF: 1:1000 |

**Directly Conjugated Antibodies**

| Antibody | Type | Source | Application |
| --- | --- | --- | --- |
| Cardiac Troponin T, Alexa Fluor® 647 | mouse monoclonal, clone 13-11 | BD Biosciences #565744 | Flow: 1:133 |
| Mouse IgG1 k Isotype Control, Alexa Fluor® 647 | mouse monoclonal, clone MOPC-21 | BD Biosciences #565571 | Flow: 1:133 |
| Nanog, Alexa Fluor® 647 | mouse monoclonal, clone N31-355 | BD Biosciences #561300 | Flow: 1:67 |
| Oct3/4, Brilliant Violet™ 510 | mouse monoclonal, clone 40/Oct-3 | BD Biosciences #563524 | Flow: 1:5 |
| Sox2, V450 | mouse monoclonal, clone O30-678 | BD Biosciences #561610 | Flow: 1:80 |

**Secondary Antibodies**

| Antibody | Type | Source | Application |
| --- | --- | --- | --- |
| goat anti-mouse IgG (H+L), Alexa Fluor® 647 | goat polyclonal | ThermoFisher #A21236 | IF: 1:500 - 1:1000 |
| goat anti-rabbit IgG (H+L),<br>Alexa Fluor® 647 | goat polyclonal | ThermoFisher #A21245 | IF: 1:500 -<br>1:1000 |
| goat anti-mouse IgG (H+L),<br>Alexa Fluor® 568 | goat polyclonal | ThermoFisher #A11031 | IF: 1:500 -<br>1:1000 |
| goat anti-rabbit IgG (H+L),<br>Alexa Fluor® 568 | goat polyclonal | ThermoFisher #A11036 | IF: 1:500 -<br>1:1000 |

### Primers and crRNA sequences (all sequences in 5’ to 3’ orientation)

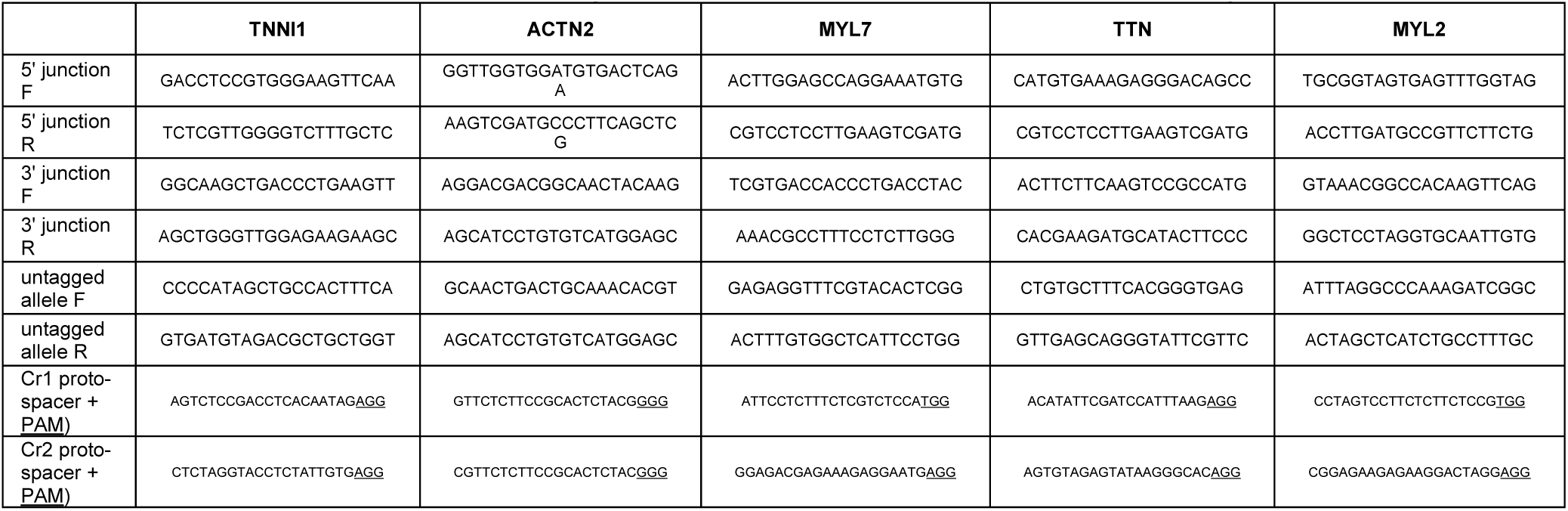

## Author Contributions

BR designed experiments, performed hiPSC editing experiments and wrote the manuscript. HM, AN and KG designed and performed cardiomyocyte differentiation experiments. JA and HM performed hiPSC and cardiomyocyte FACS experiments. MH, CH, SL, IM and RY designed and performed imaging experiments. SR designed experiments. RG designed experiments and wrote the manuscript. All authors contributed to manuscript preparation.

## Acknowledgements

We thank Colette DeLizo, Margaret Fuqua, Tanya Grancharova, Amanda Haupt, Rick Horwitz Sean Palecek, Jacqueline Smith, and Andrew Tucker for assistance in preparing the manuscript and for helpful discussions. We thank Thao Do for illustrations. The WTC line that we used to create our gene edited cell lines was provided by the Bruce R. Conklin Laboratory at the Gladstone Institute and UCSF. We thank the Allen Institute for Cell Science founder, Paul G. Allen, for his vision, encouragement, and support.

---

**Figure S3.**
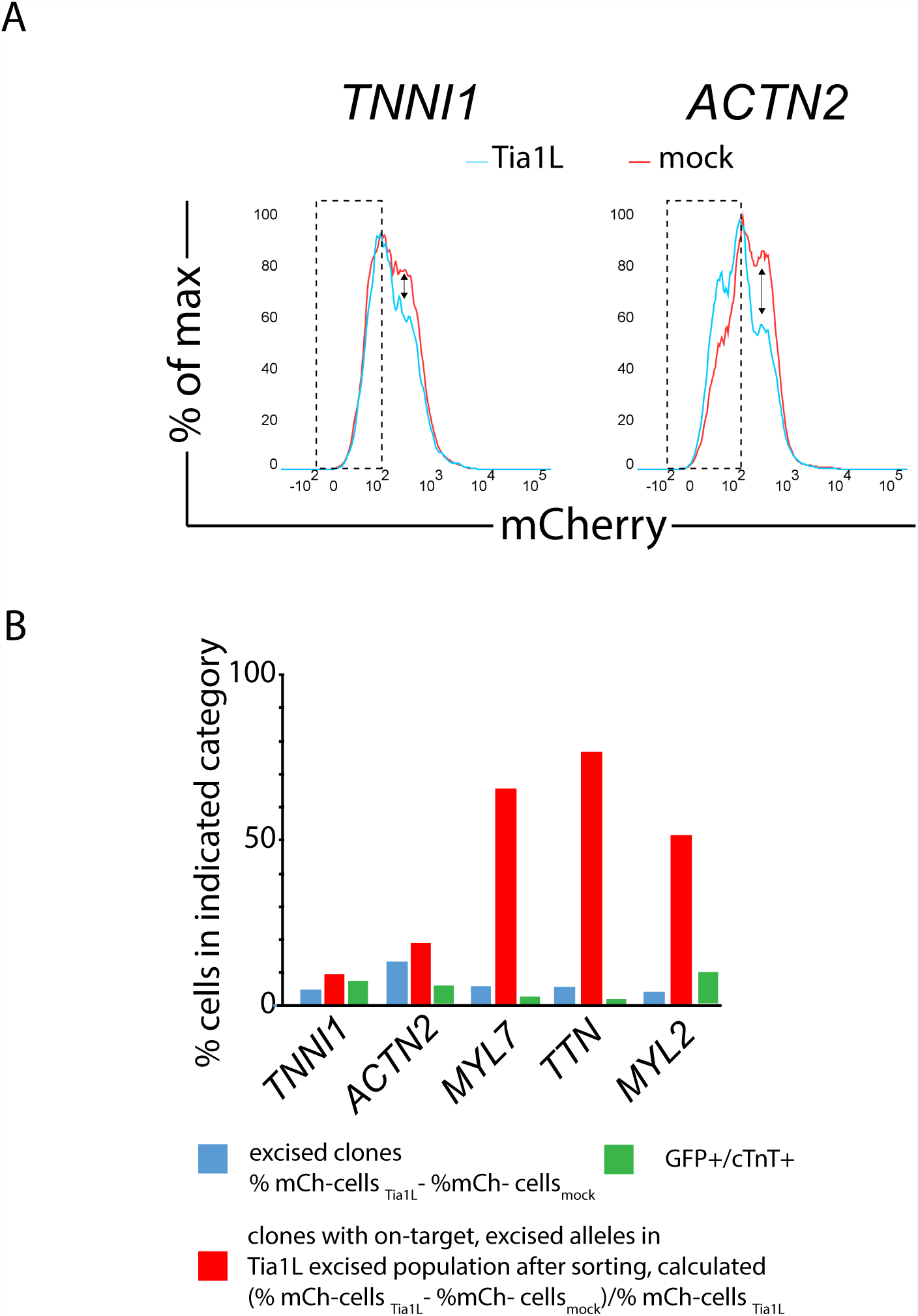
Experiments to validate tagging at *ACTN2* and *TNNI1* loci. **A.** Flow cytometry histograms display cell frequencies as a function of mCherry fluorescence from Tia1L-RNP-excised (blue) and mock-transfected (red) populations of putatively edited cells (mCherry-expressing) from the *TNNI1* and *ACTN2* targeting experiments. mCherry-negative cells were abundant in the mock condition because hPGK promoter silencing occurred. The diminished frequency of mCherry-expressing cells in Tia1L RNP-transfected populations (black arrows) was consistently observed and interpreted as evidence that excision had occurred. This contrasted with the *MYL7*, *MYL2* and *TTN* tagging experiments, where mCherry expression was more stable in non-excised cells and mCherry-negative cells were much rarer in the mock transfected condition. **B.** The estimated percentage of clones with putatively tagged alleles in the Tia1L population derived from on-target excision events was determined (red) by subtracting the percentage of mCherry-negative cells in the mock-excised population from the percentage of mCherry-negative cells in the Tia1L excised population and dividing this value by the total percentage of mCherry-negative cells in the Tia1L excised population, which were hypothesized to additionally include unexcised mCherry-negative cells with silenced promoters, mis-sorted unedited cells, and excised insertions of the donor cassette. Values calculated as the absolute excision rate in panel 3C are additionally shown, along with values obtained for the percentage of GFP+/cTnT+ cardiomyocytes in panel 3D.

**Figure 6S.**
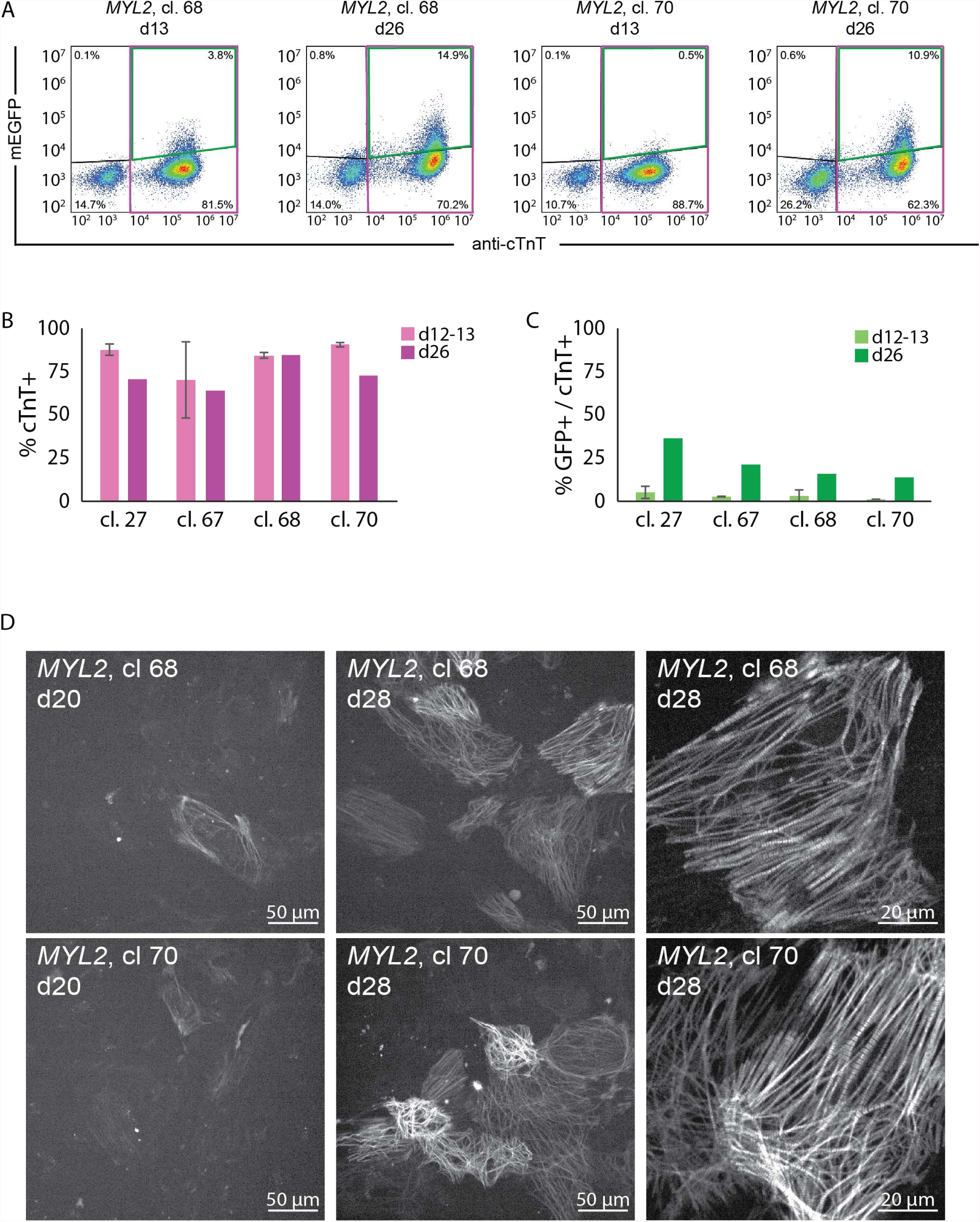
Evaluation of cardiomyocyte differentiation and mEGFP expression in *MYL2* clones. **A.** Two representative *MYL2* clones predicted to produce an in-frame endogenous mEGFP fusion protein were differentiated into cardiomyocytes and cultured for 13 and 26 days. mEGFP fluorescence in fixed cells was measured by flow cytometry (y axis) and plotted against antibody staining intensity for the cardiomyocyte marker cardiac troponin T (cTnT, x axis). Each clone displayed successful differentiation, with most cells expressing cTnT (pink box), and an increasing percentage of cardiomyocytes expressing mEGFP on day 26 compared with day 13 (cTnT+/mEGFP+, green box). Percentages of cells gated as positive and negative for both mEGFP fluorescence and cTnT staining are as indicated. **B.** The percentages of cells expressing cTnT in four precisely edited clones are shown. **C.** The percentages of cardiomyocytes (cTnT+) that were additionally expressing mEGFP are shown. In **B** and **C**, error bars are standard deviation amongst biological replicates. Day 12-13 experiments included two replicates. Day 26 experiments consisted of one replicate. **D.** Cardiomyocytes from two clones from *MYL2* editing experiments were plated on glass bottomed plates coated with PEI/laminin and were imaged live using spinning disc confocal microscopy. At day 20 (left column) and day 28 (middle column) after differentiation mEGFP expression was assessed, showing an increase in both the number of cells expressing mEGFP and the intensity of mEGFP signal. Additionally, mEGFP localization was observed specifically at sarcomeres in both clones (right column shows higher magnification). Scale bars are as indicated.

